# Vascularized midbrain assembloids show neuroinflammation and dopaminergic neuron vulnerability in Parkinson’s Disease

**DOI:** 10.1101/2025.08.06.668815

**Authors:** Anna-Sophie Zimmermann, Sonia Sabate-Soler, Alise Zagare, Kyriaki Barmpa, Kristian Haendler, Susana Rosa, Lino Ferreira, Cláudia Saraiva, Malte Spielmann, Jens Schwamborn

## Abstract

The use of micro-physiological systems has rapidly risen in the last years due to their translatability and complex cellular composition. Human midbrain-specific organoids contain neuroectoderm-derived cell types and are suitable for brain region-specific disease modeling. However, the lack of vasculature in these systems reduces oxygenation and nutrient supply. Furthermore, neurovascular interactions cannot be studied, and disease phenotypes affecting vascular and neurovascular structures cannot be assessed. To overcome these limitations, in this work, we successfully incorporated a vascular network into midbrain organoids by fusion with vascular organoids. Midbrain-vascular assembloids are enriched in vascular cells and microglia. We observed a decrease in hypoxia and cell death in these assembloids. Furthermore, microglia and endothelial cells increased their morphological complexity. Assembloids derived from a Parkinson’s disease patient carrying a LRRK2-G2019S mutation displayed a pro-inflammatory phenotype and altered electrophysiological properties. Midbrain-vascular assembloids increase the midbrain model complexity and allow for neuroinflammation studies in Parkinson’s disease.

## Introduction

Research on three-dimensional (3D) organoid models has risen over the last decades, since organoids accurately represent the cellular organization and functionality of human tissues ^1–3^. The brain is one of the most complex organs in the human body, with high cellular diversity and organization. Whole brain ^4,5^ and region-specific ^6,7^ models have been developed in the last years to recapitulate the cellular environment of the human brain. Furthermore, CNS disorders often affect specific brain areas, and region-specific models, which may be enriched in specific cell types, are key for modeling disease phenotypes that strongly impact these areas. Human midbrain-specific organoids (MOs) have been proven to be a suitable model for Parkinson’s disease (PD) since they recapitulate the two major hallmarks of the disease: neurodegeneration of dopaminergic neurons and protein aggregation ^8–10^. MOs have a heterogeneous cellular composition, including neurons, astrocytes, oligodendrocytes, and precursor cells. Neurons in MOs are functional and able to communicate with one another through synaptic contacts ^8^.

The key limitation of current MO models is the lack of mesoderm-derived cells, such as microglia and endothelial cells. Microglia have been successfully integrated into MOs, leading to lower stress levels and higher electrophysiological functionality ^11^. However, a vascularization system is still missing in MOs. The lack of oxygen and nutrients in the inner core is a common issue in 3D models, often leading to high levels of cell death ^7^. We hypothesize that the presence of a vasculature system in MOs would not only increase their resemblance to the human brain but also enhance oxygen flow in the tissue.

Blood vessels are composed of endothelial cells connected by specific junctions, pericytes, and basal lamina containing smooth muscle cells ^12,13^. In the brain, they interact with neuronal projections, astrocyte feet, and microglia, forming the neurovascular unit ^14–16^. Recent work has shown that engineered scaffolds mimicking vascular diffusion can enhance MO maturation and reduce hypoxia ^17^. However these systems rely on artificial scaffolds and have not been explored in the context of PD. Moreover, neurovascular interactions in human MOs remain largely unexplored. We hypothesize that, in a vascularized 3D model of the human brain, vascular cells establish connections with neurons, astrocytes, and microglia, that may have a positive impact on microglia morphology or functionality. Beyond improving oxygenation, we also speculate that this complex 3D system is suitable for PD disease modeling, and cellular phenotypes related to neuroinflammation may be enhanced in this advanced model.

In this article, we integrate a vasculature system in MOs by co-culture of midbrain and vascular organoids. Our findings show that hypoxia levels in midbrain-vascular assembloids (MOVOs) are indeed reduced in comparison to MOs, suggesting an enhanced oxygen supply. Furthermore, by co-culture of assembloids with macrophage precursors and differentiation into microglia in 3D, we obtained a complex assembloid system (MGL-MOVOs) where microglia spatially interact with endothelial cells and neuron processes, mimicking the neurovascular unit *in vitro*. Interestingly, microglia and endothelial cells in MGL-MOVOs display a more complex morphology compared to MOs. In a PD MGL-MOVO model, derived from patients’ iPSCs carrying a LRRK2-G2019S mutation, we observed altered electrophysiological properties as well as a pro-inflammatory phenotype. Furthermore, single nuclei RNA sequencing (snRNA-Seq) data highlighted a specific dopaminergic neuron cluster in mutant assembloids, characterized by enhanced expression of ferroptosis and cell death-related genes. The vascularized, microglia-containing midbrain organoid model presented here enables the assessment of disease-relevant phenotypes, including neurovascular interactions, neuroinflammatory alterations, and dopaminergic neurons vulnerability.

## Results

### MOVOs express specific vascular markers and show reduced hypoxia levels

To integrate a vascular system into midbrain organoids (MOs, Figure 1A), we derived neuro-epithelial stem cells (NESCs) from two wildtype, healthy control (HC) iPSC lines (HC2 and HC1) and differentiated them into MOs. In parallel, we generated vascular organoids following a published procedure ^18^. Then, we co-embedded them in an extracellular matrix gel with cell line-matched MOs ^19^. We cultured the vascularized midbrain organoids (MOVOs) for 24 more days, until day 30 of differentiation (Figure 1A). MOs were used as control. By day 30 of differentiation, MOVOs from lines HC1 and HC2 showed the presence of the endothelial cell marker CD31, the dopaminergic neuron marker TH, and the pan-neuronal marker MAP2 (Figure 1B, C). Immunofluorescence staining for the basement membrane and extracellular matrix protein Collagen IV showed positive structures in assembloids, which organized spatially like CD31-positive endothelial cells (Figure 1B). Staining for alpha smooth muscle actin (αSMA), a capillary pericyte and vascular smooth muscle cell marker showed positive structures (Figure 1B). The presence of vascular-specific adherens junctions between endothelial cells was confirmed by VE-Cadherin staining in assembloids (Figure 1B). High-magnification immunofluorescence images showed the organization of CD31-positive endothelial cells in a tubular-like manner (Figure 1D). Overall, we show the presence of endothelial cells connected by tight junctions, pericytes, and basal lamina in MOVOs, which seemed to be organized in vessel-like structures.

**Figure 1.**
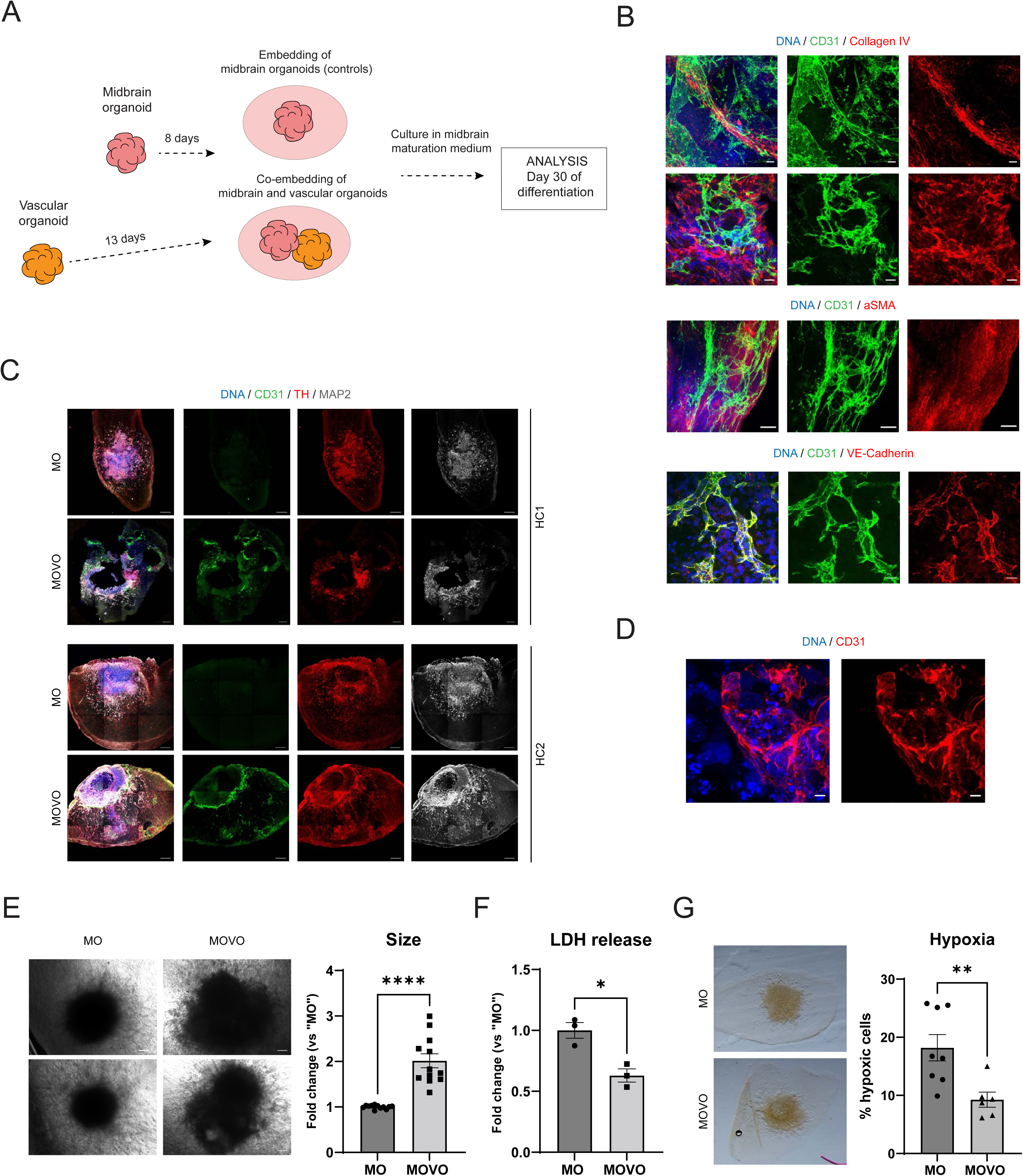
MOVOs include blood-vessel markers and show reduced LDH and hypoxia levels. **A.** Schematic representation of the organoid co-culture and generation of MOVOs. **B.** Immunofluorescence staining showing CD31 and Collagen IV in assembloids (scale bar = 50 µm in upper panels, 20 µm in bottom panels, line HC2, 70 µm sections). Staining for CD31 and alpha smooth muscle actin (αSMA) in an MOVOs (scale bar = 50 µm, line HC2, 70 µm sections). Staining for CD31 and VE-Cadherin in MOVOs (scale bar = 20 µm, line HC2, 70 µm sections). **C.** Immunofluorescence images in midbrain MOs and MOVOs from lines HC2 and HC1, showing the presence of CD31, TH and MAP2 at day 30 of differentiation (scale bar = 200 µm, 70 µm sections for line HC2 and 120 µm for line HC1). **D.** High-magnification immunofluorescence image showing a CD31-positive structure with tubular morphology in a MOVO (scale bar = 5 µm, line HC2, 70 µm sections). **E.** Representative bright field images showing MOs (left) and MOVOs (right) from lines HC2 (upper panels) and HC1 (bottom panels) at day 30 of differentiation. Bar graph representing MO and MOVO area at day 30 of differentiation (each dot represents an organoid or assembloid, n = 6 (2 cell lines (HC2 and HC1), 3 batches). Error bars = SEM). **F.** LDH release normalized to size in MOs and MOVOs from line HC1 at day 30 of differentiation (each dot represents a pooled supernatant sample from 3 organoids or assembloids from one batch, n = 3, 3 batches. Error bars = SEM). **G.** Representative images showing the Hypoxyprobe staining in an MO (up) and a MOVO (bottom, left panels). Bar graph representing the percentage of hypoxic area in MOs and MOVOs at day 15 of differentiation (each dot represents an organoid (1-3 organoids used per condition, n= 4, 2 cell lines, 2 batches). Error bars = SEM, right panel). *p < 0.05, **p<0.01, ***p<0.001, ****p<0.0001 using an unpaired t-test in A, B, and C.

After confirming the presence of a vascular network in MOVOs, we assessed its effects on cell viability and hypoxia. First, after measuring the MO and MOVO area at day 30 of differentiation, we confirmed that MOVOs are significantly larger than MOs (Figure 1E). We performed a viability LDH assay and observed a significantly lower LDH release in MOVOs compared to MOs (Figure 1F), indicating that MOVOs have lower levels of cytotoxicity. To understand if the lower cytotoxicity level was related to a better oxygenation of the organoids, we measured hypoxia levels in MOs and MOVOs using a Hypoxyprobe assay and staining. MOVOs showed a significantly lower number of hypoxic cells, suggesting that the vascular system may participate in decreasing hypoxia in MOVOs (Figure 1G).

### Cellular diversity of MGL-MOVOs

After studying the effects of the vascular network in MOVOs, we investigated the neurovascular interactions in the system. For this, we integrated macrophage precursor cells into midbrain organoids (resulting in MGL-MOs, midbrain organoids with microglia) or into vascularized midbrain organoids (resulting in MGL-MOVOs, vascularized midbrain organoids with microglia) from cell line HC1 (Figure 2A). We observed macrophage precursor integration into the system (Figure 2B). After sectioning and staining, we detected IBA1-positive cells in MGL-MOs and MGL-MOVOs, and CD31-positive cells exclusively in MGL-MOVOs (Figure 2C). These cells formed a vascular network characterized by the presence of the tight junction marker Occludin, the endothelial and astrocyte-associated channel Aquaporin-4, and the pericyte marker PDGFRβ. The results revealed co-localization of these structures within the MGL-MOVOs (Figure 2D). To gain deeper insights into the cellular diversity in MGL-MOVOs, as well as the features of the microglia and vascular compartments, we performed single-nuclei RNA sequencing (snRNA-Seq). Clustering analysis revealed the exclusive presence of microglia and vascular-associated populations (corresponding to Endothelial cells 1, 2 and 3, Mesoderm-like cells, and Megakaryocyte precursor cells) in MGL-MOVOs, all absent in MOs (Figure 2E). Approximately 52% of cells in MOs and 12% in MGL-MOVOs exhibited neuronal characteristics (corresponding to clusters of Neurons, Cholinergic neurons, and Dopaminergic neurons 1). In contrast, vascular-associated cells made up over 65% of the MGL-MOVO composition. This shift aligns with published single-cell RNA-Seq data from postmortem human midbrain tissue, which revealed a cellular landscape composed of glial, vascular, microglial, and multiple neuronal subtypes, with neurons comprising only 10-20% of total cells and dopaminergic neurons representing just 1-5%, even within the substantia nigra ^20,21^. To confirm our observation, we saw that vascular-associated cells positive for the collagen markers *COL1A1* and *COL5A1* are highly represented in MGL-MOVOs but not in MOs (Figure S1A). Additionally, glial cells, including astrocytes and microglia, accounted for 3.80% of the total MGL-MOVO cell population (Figure 2F and Table S3). Differential gene expression (DEG) analysis across all cell types, followed by process network enrichment, revealed significant enrichment in networks related to development and neurogenesis, cell adhesion and synaptic contact, and cell-matrix interactions in MGL-MOVOs compared to MOs (Figure S1B). These findings suggest that MGL-MOVOs exhibit a more advanced state of cellular maturation. In line with this, expression of neurotrophic factors (*BDNF*, *GDNF*, *NGF*, *NTF3*) was elevated in MGL-MOVOs (Figure S1C), as was the expression of key synaptic genes including *NLGN1*, *RIMS1*, *PCLO*, and *SYN3* (Figure S1D). These findings suggest that the presence of microglia and a vascular component fosters an environment that supports enhanced neuronal maturation, neurotrophic signaling, and synaptic development, closely reflecting features of the human midbrain *in vivo*, prompting us to investigate whether microglial morphology similarly mirrors this more mature and complex microenvironment.

**Figure 2.**
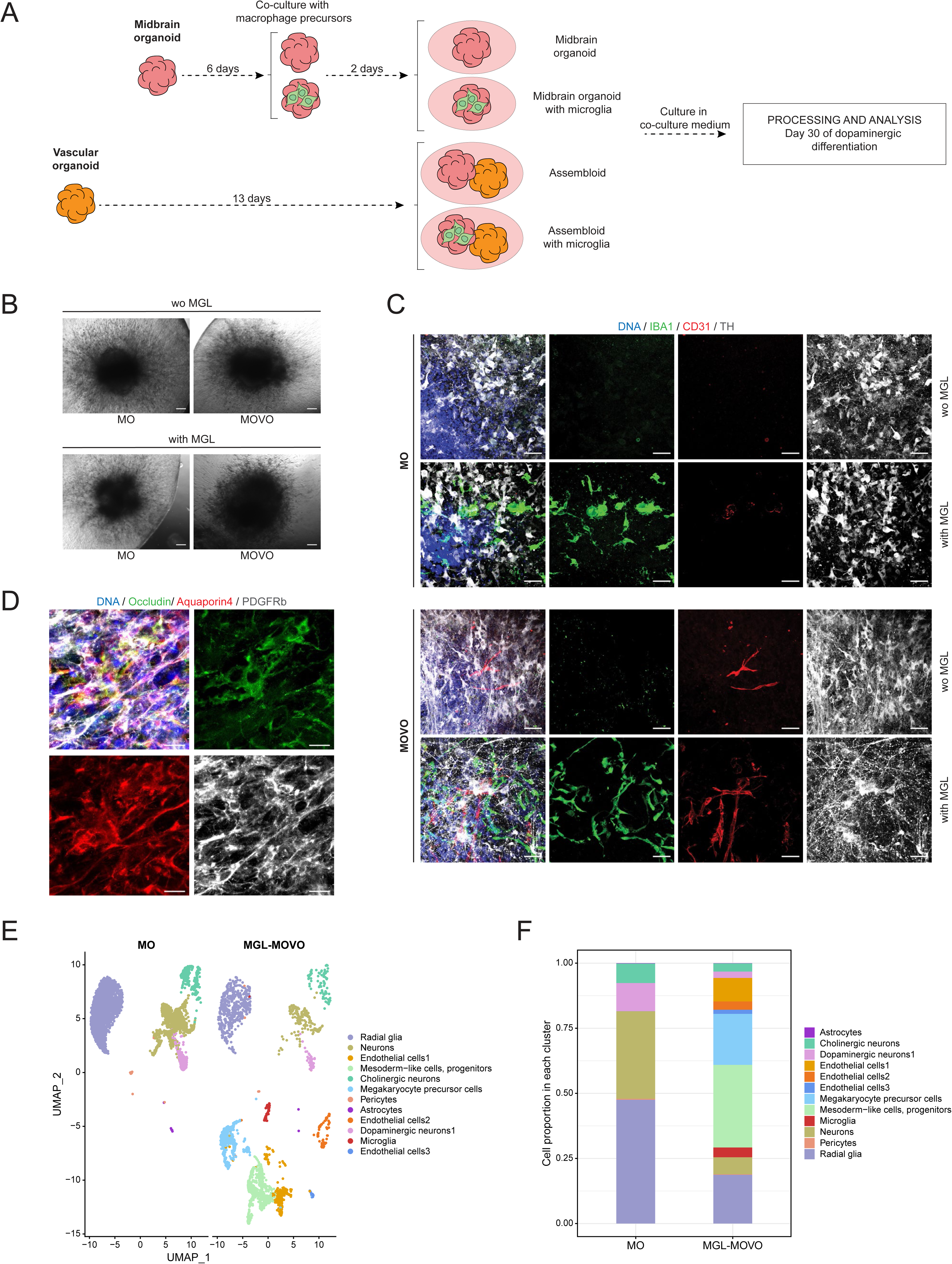
MGL-MOVOs are enriched in resting microglia and cellular components of blood vessels. **A.** Schematic representation of the co-culture procedure for the integration of microglia into MOs and MOVOs. **B.** Bright field images showing MOs and MOVOs co-cultured with microglia (MGL) and their respective controls (Scale bar = 200 µm). **C.** Immunofluorescence images of MOs and MOVOs without and with MGL, for the markers IBA1, CD31, and TH (Scale bar = 50 µm. Images correspond to an orthogonal projection of planes). **D.** Immunofluorescence image of an MGL-MOVO section stained for occludin, aquaporin and PDGFRβ (Scale bar = 10 µm. Images correspond to an orthogonal projection of planes). **E.** UMAP visualization of snRNA-Seq data revealed six distinct cell clusters in MOs and 12 distinct cell clusters in MGL-MOVOs. Each dot represents a single cell and is coloured according to the cell identity. **F.** Proportions of different cell types in HC1 MOs and MGL-MOVOs.

### Microglia increase their morphological complexity in MGL-MOVOs

Using 3D reconstructions of immunostaining images, we observed close interaction between microglia and endothelial cells in MGL-MOVOs (Figure 3A, left panel). Furthermore, this microglia-endothelial complex made contact with TH-positive projections, showing neurovascular unit-like interactions (Figure 3A, right panel). Microglia and endothelial cells in MGL-MOVOs seemed to have a more ramified morphology compared to MGL-MOs or MOVOs, respectively. These observations were confirmed with image analysis, where we measured the length sum of processes per cell. The significantly increased length of microglia processes observed in MGL-MOVOs compared to MGL-MOs (Figure 3B) is characteristic of surveillant microglia in a homeostatic state, in contrast to the ameboid morphology typically exhibited by activated microglia ^22^. Furthermore, the length of endothelial cell processes was higher in MGL-MOVOs (Figure 3C). Thus, in terms of morphology and process length, both microglia and endothelial cells seemed to benefit from each other’s presence in the system. Confocal imaging showed that endothelial cells expressing CD31 and CD34 formed interconnected networks (Figure 3D). 3D reconstructions further validated the presence of self-organizing vascular networks composed of lumen-forming endothelial cells (CD31 and CD34) (Figure 3E), validating CD34 as a reliable marker for visualizing endothelial cells in MGL-MOVOs alongside CD31. To investigate vascular structures and neurovascular unit-like interactions in MGL-MOVOs in a more advanced developmental stage, MGL-MOVOs were cultured for 120 days of differentiation. This extended maturation period allowed for the assessment of more developed neurovascular interactions within the system. Immunofluorescence staining and 3D reconstruction for CD34, IBA1, and the astrocyte marker glial-fibrillary acidic protein (GFAP) showed the close association and interaction between microglia, astrocytes, and endothelial cells (Figure 3F, 3G, and Supplementary video). In addition, a co-localization of PDGFRβ, CD34 and COL1A1 was detected along vessel-like structures (Figure S1E), indicating the formation of perivascular interactions.

**Figure 3.**
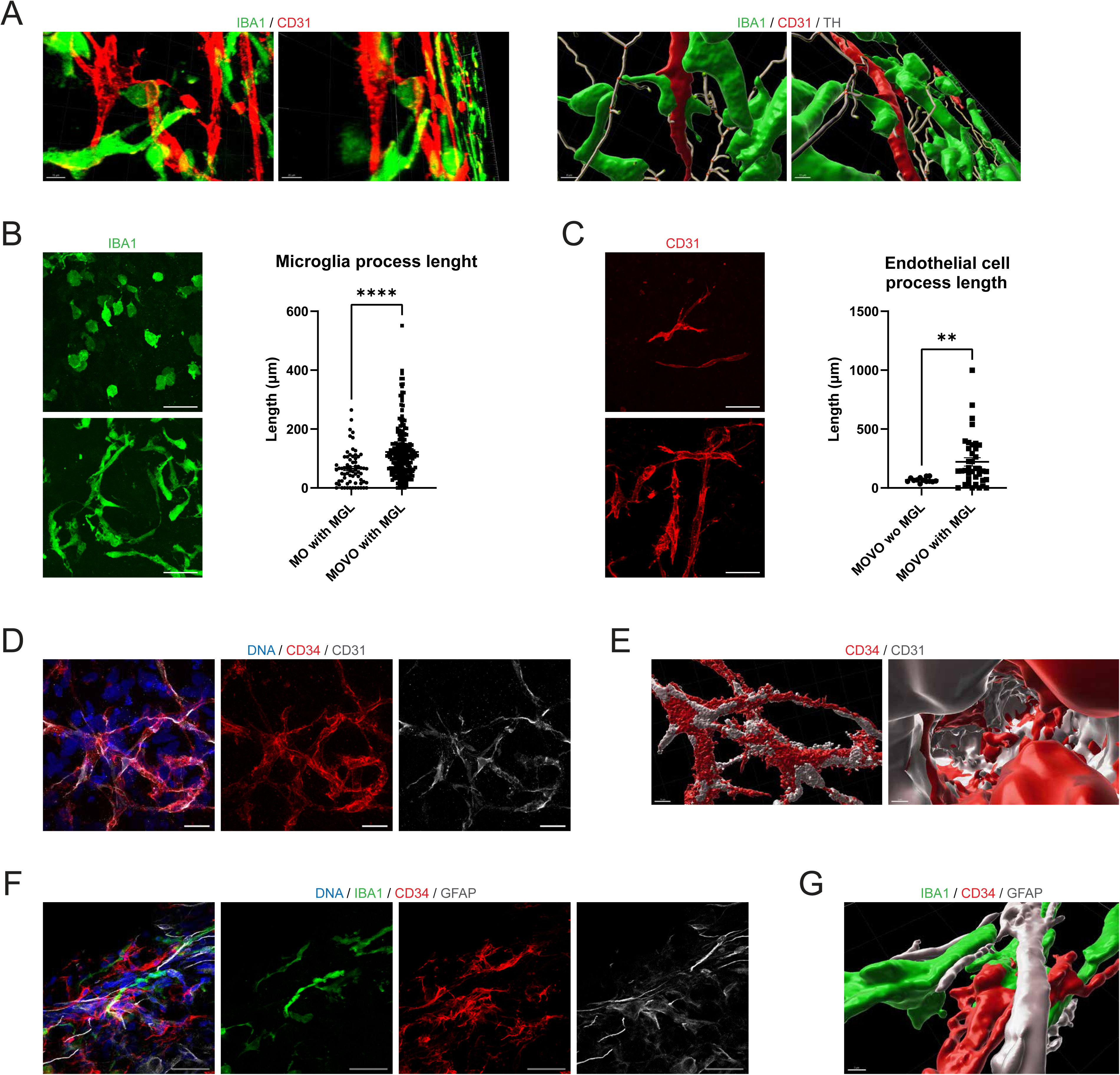
Microglia and endothelial cells increase their morphological complexity in MGL-MOVOs and interact in a neurovascular unit-like manner. **A.** 3D reconstruction of the interaction of endothelial cells (CD31) with microglia (IBA1) from different angles (left panels, scale bar = 15 µm in left panel and 20 µm in right panel). 3D reconstruction representing the interaction of endothelial cells, microglia, and dopaminergic neurons (TH, filaments on image) from different angles (right panels, scale bar = 8 µm in left panel and 10 µm in right panel) in MGL-MOVOs. **B.** Representative immunofluorescence images of an MO with microglia (MGL, up) and an MGL-MOVO (bottom, left panels) stained for IBA1 (Scale bar = 50 µm). Bar graph showing a significant increase in microglia process length. **C.** Representative immunofluorescence images of a MOVO (up) and an MGL-MOVOs (bottom, left panels) stained for CD31 (Scale bar = 50 µm). Bar graph showing a significant increase in endothelial cell process length. **D.** Immunofluorescence images of CD31 and CD34 expressing endothelial cells show the establishment of vascular networks within MGL-MOVOs at 30 days of differentiation (scale bar = 50 µm). **E.** 3D reconstruction of vascular network organization (CD31 and CD34) in MGL-MOVOs at 30 days of differentiation (scale bar = 10 µm in left panel and 5 µm in right panel). **F.** Immunofluorescence images of an MGL-MOVO stained for IBA1, CD34, and GFAP at 120 days of differentiation (Scale bar = 50 µm). **G.** 3D reconstruction representing the interaction of endothelial cells, microglia, and astrocytes in MGL-MOVOs at 120 days of differentiation (Scale bar = 2 µm). In B and C, each dot represents one cell. The length is calculated as a sum of all processes in a cell (2 organoids used per condition, n= 2, 1 cell line, 2 batches. Error bars = SEM). *p < 0.05, **p<0.01, ***p<0.001, ****p<0.0001 using a Mann-Whitney test.

### Differential gene expression analysis reveals dopaminergic neuron vulnerability in LRRK2-PD MGL-MOVOs

Having characterized and confirmed neurovascular-like interactions in HC MGL-MOVOs, we next wanted to explore their potential for Parkinson’s disease modeling. Accordingly, we compared the cellular compositions in PD MGL-MOVOs (derived from patient iPSCs with the LRRK2-G2019S mutation) and MGL-MOVOs from a healthy control individual. Comparison was done via visualization in UMAP embeddings (Figure 4A). Clusters corresponding to radial glia, neurons, endothelial cells (1, 2, and 3), mesoderm-like cells, cholinergic neurons, megakaryocyte precursor cells, pericytes, astrocytes, dopaminergic neuron 1 (DNs1), and microglia were detected in both MGL-MOVO types, indicating a substantial overlap in cellular composition. However, the dopaminergic neuron 2 (DNs2) cluster was exclusively present in PD MGL-MOVOs. The presence of two distinct dopaminergic neuron clusters (DNs1 and DNs2), with DNs2 specifically emerging in PD MGL-MOVOs, suggests that this cluster may differentially contribute to neurodegenerative processes. To identify the key differences between DNs1 and DNs2 clusters, we performed a DEG analysis across HC and PD MGL-MOVOs, followed by a process network enrichment analysis. This analysis revealed a significant enrichment in process networks associated with cell adhesion and synaptic contacts (Figure 4C). To further explore the distinct genes expression profiles of DNs1 and DNs2, we visualized the fold changes of genes involved in the top significantly enriched process network: cell adhesion and synaptic contact (Figure 4D). Notably, genes linked to synaptic adhesion and scaffolding (*NRXN1*, *NRXN2*, *NLGN1*, *LSAMP*, *NTM*, *OPCML*, *CNTN1*, *SHANK2*), synaptic transmission (*GRIA1*, *GRIA4*, *GRIK2*, *GRIN2A*, *GRIN2B*, *GRIP1*, *CAMK2A, CAMK2B*), and axonal guidance (*EPHB1*) were found to be downregulated in DNs2, suggesting functional differences between these two dopaminergic neuron subpopulations. Further supporting this distinction, a dot plot visualization revealed a reduced expression of critical markers of dopaminergic neuron identity and midbrain specification (*KCNJ6*, *LMX1B*, and *EN1*) in DNs2. This transcriptional reduction may reflect a developmentally altered phenotype in DNs2. Additionally, the plot highlighted an increased expression of ferroptosis-related factor *GPX4* and apoptosis-related factor *CASP7* in DNs2 compared to DNs1 (Figure 4E). The elevated *GPX4* levels may indicate an adaptive cellular response, while the higher *CASP7* expression suggests a heightened state of vulnerability in DNs2.

**Figure 4.**
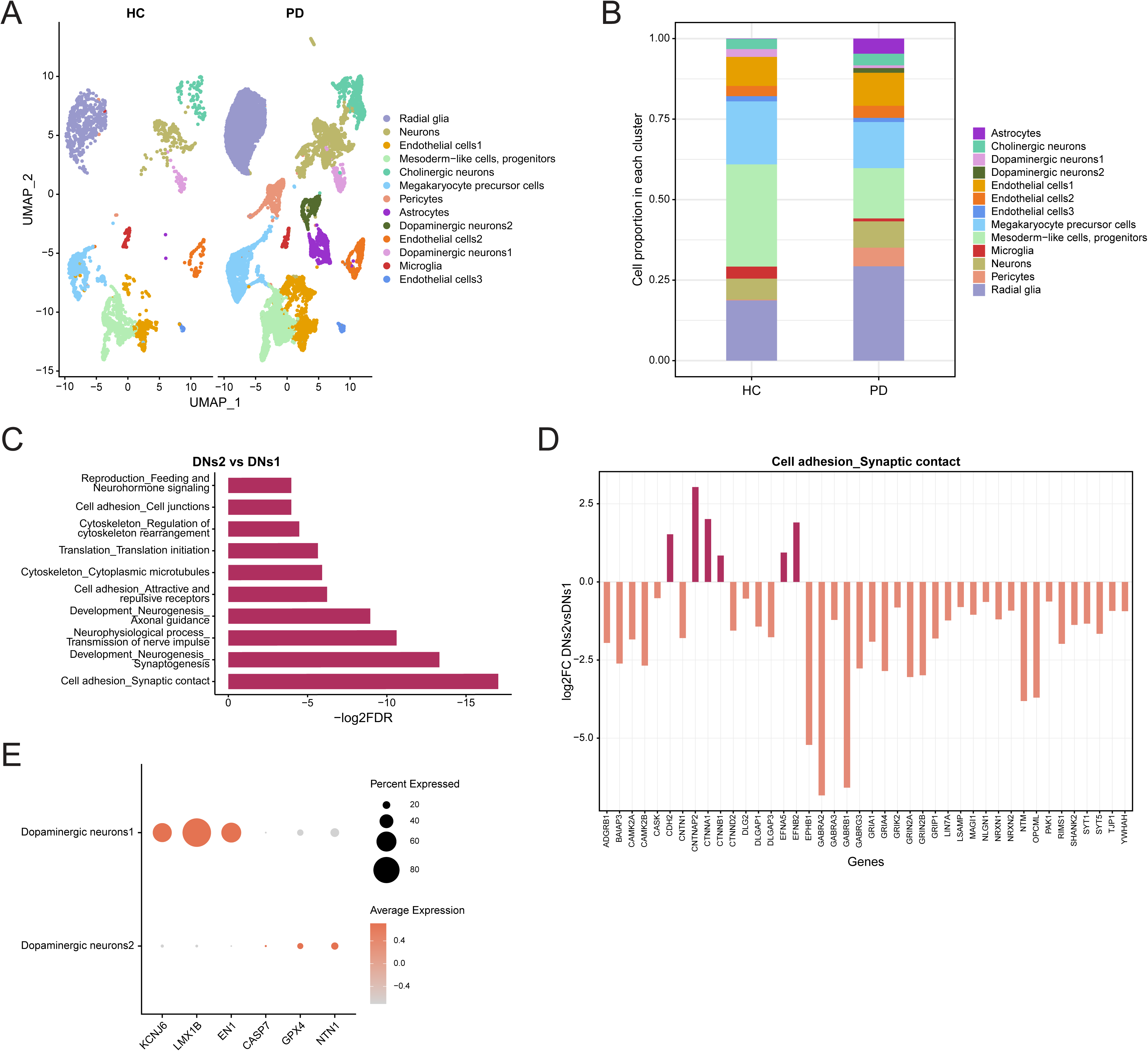
LRRK2 PD MGL-MOVOs show dopaminergic neuron vulnerability. **A.** UMAP of cell clusters in HC1 and LRRK2 G2019S PD (PD) MGL-MOVOs. Each dot represents a single cell and is coloured according to the cell identity. **B.** Proportions of specific cell abundance in different cell types in HC1 and PD MGL-MOVOs. **C.** Bar graph representing the significantly enriched process networks based on the differential gene expression analysis between dopaminergic neurons 1 (DNs1) and dopaminergic neurons 2 (DNs2) in MGL-MOVOs (p adj. value <0.05, -log2FDR threshold >0.25). **D.** Fold changes of genes selected from the top enriched pathway between DNs1 and DNs2 in MGL-MOVOs (p adj. value <0.05, -log2FDR threshold >0.25). **E.** Dot plot illustrating differential gene expression of 6 key genes in DNs1 and DNs2.

Further, we investigated the electrophysiological activity of both HC and PD assembloids. We recorded spontaneous electrophysiological activity using a multi-electrode array (MEA) system (Figure S2A). Our analysis revealed that HC MGL-MOVOs exhibited a significantly higher mean firing rate (MFR) at days 30 and 40 of differentiation (Figure S2B). Moreover, throughout the recording period from day 25 to 40, HC MGL-MOVOs consistently maintained a higher MFR (Figure S2C) and HC MGL-MOVOs demonstrated a significantly higher number of spikes than PD MGL-MOVOs at days 30 and 40 (Figure S2D), further indicating LRRK2-G2019S-mutant-associated impaired neuronal functionality.

### LRRK2-PD MGL-MOVOs show altered microglia morphology and neuroinflammation phenotypes

To further explore the effects of the LRRK2-G2019S mutation, we analysed microglia ramifications in HC and PD MGL-MOVOs. Microglia in PD MGL-MOVOs appeared to exhibit a less ramified morphology compared to those in HC MGL-MOVOs (Figure 5A). This observation was quantitatively confirmed through image analysis. The results demonstrated significantly longer microglial processes in HC MGL-MOVOs compared to PD (Figure 5B). To gain deeper insights into microglial phenotypes, we examined cell-type-specific gene expression in microglia from HC and PD MGL-MOVOs through snRNA-Seq (Figure 5C). A heatmap displaying microglia-specific marker expression revealed distinct expression patterns across HC and PD MGL-MOVOs. Dot plot visualizations highlighted two groups of differentially expressed genes. The first group (Figure 5D) included genes involved in immune modulation and cytokine signaling (*IL7*, *TXLNA*, *IL15*, *IL4*, *IL5*, *IL10*), which were more highly expressed in HC MGL-MOVOs, suggesting a transcriptional profile associated with immune regulatory capacity. The second group (Figure 5E) included genes related to microglial environment sensing, phagocytosis, and lipid metabolism (*LGALS3*, *AXL*, *ITGAX*, *SPP1*, *CCL2*, *CSF1*, *LPL*, *TREM2*). Notably, triggering receptor expressed on myeloid cells 2 (*TREM2*), a key regulator of microglial activation and homeostasis ^23–25^, exhibited higher expression in HC compared to PD (Figure 5E), suggesting a more active microglial state in HC MGL-MOVOs. Since *TREM2* plays a crucial role in phagocytosis and immune regulation, its reduced expression in PD MGL-MOVOs may indicate an altered microglial response in the context of the LRRK2-G2019S mutation. Since microglia dynamically respond to environmental cues by producing both anti- and pro-inflammatory cytokines, the latter of which, such as IL-1, IL-6, INF, and TNF-α, have been implicated in PD pathogenesis ^26^. Flow cytometry analysis further confirmed a significant upregulation of TNF-α, IL-1β, INFγ, and IL-6 in PD MGL-MOVOs (Figure 5F). Collectively, these findings suggest that the LRRK2-G2019S mutation alters microglial function and enhances inflammatory activity.

**Figure 5.**
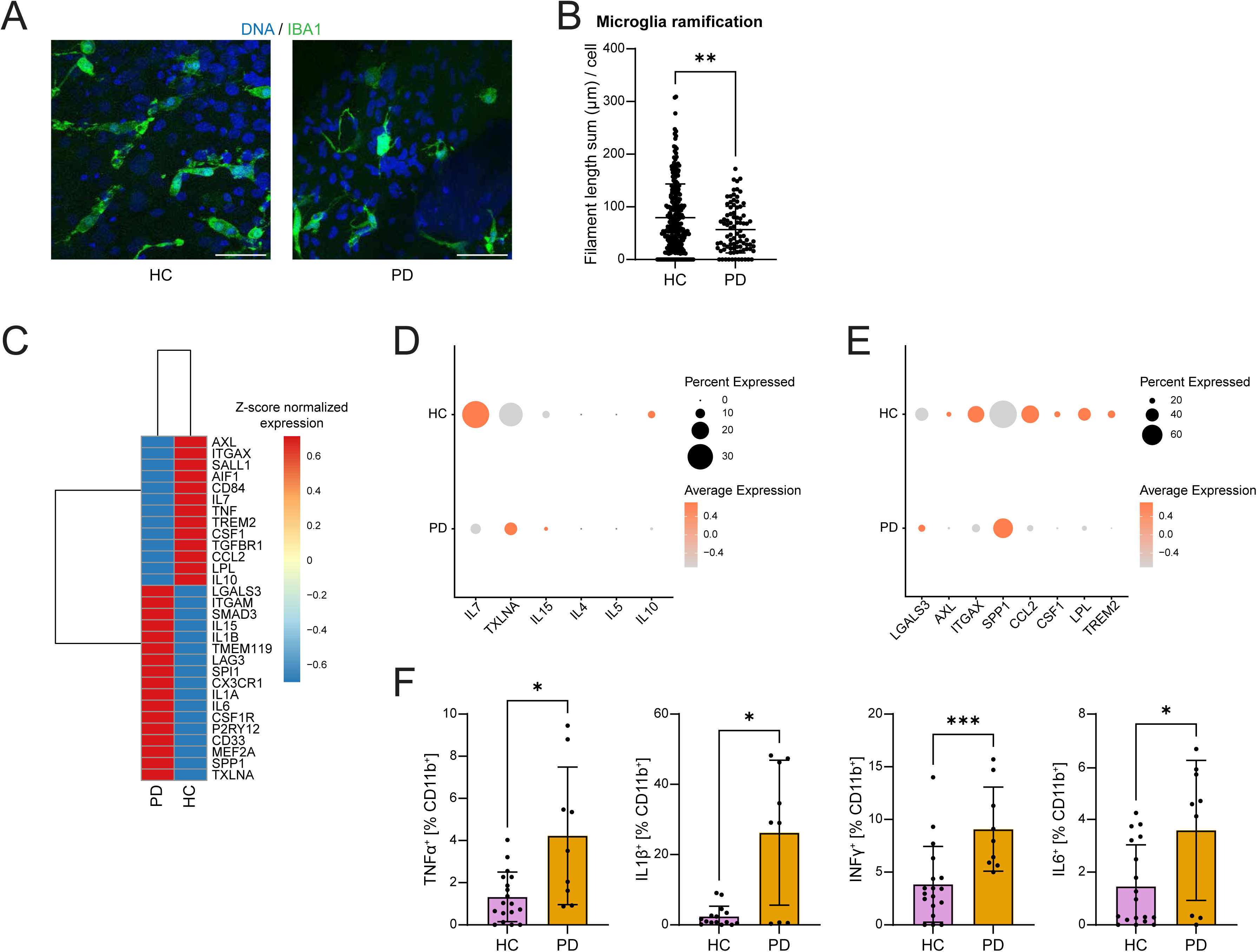
Microglia decrease their ramification and exacerbate their pro-inflammatory profile in LRRK2 PD MGL-MOVOs. **A.** Immunofluorescence images of MGL-MOVO sections from HC and PD MGL-MOVOs for IBA1 (Scale bar = 50 µm). **B.** Graph showing significantly lower microglia ramification in PD compared to HC1 MGL-MOVOs. Each dot represents one cell. The length is calculated as a sum of all processes in a cell (1 MGL-MOVO used per condition, n = 3, 3 cell lines, 3 batches. Error bars = SD). *p < 0.05, **p<0.01, ***p<0.001, ****p<0.0001 using a Mann-Whitney test. **C.** Heatmap showing microglia-specific marker expression in the microglia cluster of HC1 and PD MGL-MOVOs. **D.** Dot plot showing microglial genes involved in immune modulation. **E.** Dot plot showing microglial genes linked to environmental sensing, phagocytosis, and lipid metabolism. **F.** Flow cytometry results showing a significant increase in the pro-inflammatory cytokines Tumor necrosis factor alpha (TNFα), Interleukin-1 beta (IL1β), Interferon-gamma (INFγ), and Interleukin-6 (IL-6) in microglia from PD MGL-MOVOs. Each dot represents one measurement (n = 4 pooled MGL-MOVOs per cell line, 4 cell lines, 3 batches, 2-3 technical replicates. Error bars = SD). *p < 0.05, **p<0.01, ***p<0.001, ****p<0.0001 using a Mann-Whitney test.

In summary, the vascularized, microglia-containing midbrain assembloid model recapitulates key features of human midbrain architecture and cellular diversity. This model reveals disease-relevant phenotypes in the context of Parkinson’s disease, including dopaminergic vulnerability and altered microglial responses.

## Discussion

Integration of a vasculature system in whole brain and region-specific organoids has been attempted over the last decades ^27–29^. Amongst the approaches that have been assessed is the transplantation of organoids into mice to promote the angiogenesis of the organoid tissue ^30^. This has been done with a prior co-culture with endothelial progenitor cells ^27,31^, and mature endothelial cells ^28^. However, the need for transplantation into animal models represents some limitations in terms of reproducibility, cost, and ethical concerns. Various protocols have been explored to introduce vasculature into brain organoids, including the external incorporation of mesoderm-derived cells ^27,32^ and the integration of artificial scaffolds designed to mimic a circulatory system in organoids ^29,33^. Cerebral organoids have been co-cultured with vascular organoids ^34,35^. However, these approaches often rely on transplantation or fail to generate stem cell-based vascularized, region-specific organoids, particularly for the midbrain. In this work, we successfully co-cultured vascular organoids with MOs. MOVOs were rich in endothelial cells, a crucial component of blood vessels ^13^. Collagen-positive staining in MOVOs indicated that basal lamina-like structures are formed. CD31 seemed to spatially organize with VE-cadherin, indicating that endothelial cells are connected through adherens junctions, crucial to maintaining vascular integrity ^36^.

In 3D cell culture models, high cell death levels are often observed in the inner core of the tissue. This is mainly caused by the lack of oxygen and nutrients, leading to high hypoxia levels in the core ^7,37–39^. Some attempts to overcome this issue include protocol optimisation to decrease organoid size and, therefore, allow nutrient and oxygen diffusion to the organoid tissue ^19^. Other approaches focused on the slicing of live organoids and culturing organoid sections to enhance oxygen and nutrient diffusion ^40^. Our findings of reduced hypoxia levels and cell death in MOVOs align with previous studies assessing cell death and hypoxia in vascularized cortical organoids; a lower coverage of hypoxia-inducible factor 1-α (HIF-1α)-positive regions has been observed in *ETV2*-overexpressing vascularized cortical organoids compared to controls ^41^. The lower levels of hypoxia and cell death in MOVOs suggest that the presence of blood vessel-like structures may result in a more efficient organoid oxygenation and nutrient supply. In addition to enhancing oxygen and nutrient supply, endothelial cells are known to secrete trophic factors that support the survival, maturation, and maintenance of surrounding neural cells, including neurons and glia. Heterotypic interactions between endothelial cells and neural components are thus likely to contribute to tissue health in MOVOs ^42^. The integration of a vascular network may result in an overall less compact tissue, allowing for oxygen to flow through the structure.

Previous studies have focused on the cellular interactions between microglia and endothelial cells in the neurovascular unit. Some of them have used animal models to assess the spatial interactions and contacts between cells ^43^ and cell signalling between the microglia and endothelium compartments ^44^. This has been assessed in healthy conditions but also in disease ^45,46^ and aging ^47^. Apart from animal models, organ-on-chip technologies have been applied to mimic a neurovascular unit on-chip ^48,49^. However, little is known about cellular and molecular interactions in the neurovascular unit in iPSC-derived human brain organoids. To study neurovascular interactions in 3D, we successfully integrated microglia in MOVOs. Immunofluorescence images and 3D reconstructions showed that microglia made contact with endothelial cells and TH-positive neuron projections and GFAP-positive astrocyte feet, suggesting neurovascular-like interactions in MGL-MOVOs. Notably, microglial exhibited increased ramification in the presence of vasculature, a morphology typically associated with homeostatic surveillance and functional maturity ^50,51^. Furthermore, there is a positive correlation between microglia elongation and ramification and developmental age ^52,53^. This indicates that microglia in assembloids could fit a healthy and mature profile. Assessing cytokine and chemokine secretion, as well as microglial maturation, would be of high value to decipher the molecular implications of these morphological changes. Similarly, endothelial cells also extended more elaborate processes, suggesting a bidirectional enhancement of structural complexity. Endothelial cells undergo an elongation and sprouting before vessel formation and closure ^54^. Thus, the higher elongation of endothelial cells in assembloids with microglia compared to the ones without microglia could indicate that the process of tube formation could be enhanced. Further analysis of later time points, where vessel formation may be advanced, would be of high value to confirm this hypothesis. Furthermore, studies have shown that elongated endothelial cells suppress the activation of monocytes *in vitro* ^55^. This suggests that factors in the endothelial secretome may contribute to the suppression of immune cell activation. Oppositely, dysfunction of endothelial cells in pathological conditions such as stroke and ischemic injury leads to microglial activation, where microglia secrete pro-inflammatory cytokines and change their morphology into an amoeboid shape ^56,57^. Studying the effects of the vascular secretome on microglia in MGL-MOVOs would be of high interest to elucidate the mechanisms behind the morphological changes of microglia..

Mutations in leucine-rich repeat kinase 2 (LRRK2) are the most common genetic form of PD. The *LRRK2* G2019S is the most prevalent point mutation in LRRK2-PD ^58,59^. Previous studies using iPSC-derived dopaminergic neurons and midbrain organoids have reported impaired neuronal differentiation, reduced morphological complexity, and dysregulation of synaptic and cytoskeletal pathways in LRRK2-PD models ^8,60–63^. However, the LRRK2-PD pathophysiology has not been studied in advanced human brain assembloid systems. Given the growing evidence of neurovascular dysfunction in PD, modeling the disease in a vascularized immunocompetent system may offer additional insights beyond traditional neuron-centric models. Vascular alterations such as blood-brain barrier disruption, endothelial inflammation, and impaired neurovascular coupling have been implicated in PD pathophysiology and may contribute to both neurodegeneration and glial dysregulation ^64–66^. In this study, snRNA-Seq analysis of MGL-MOVOs revealed greater cellular diversity, including endothelial cells, microglia, and megakaryocyte-precursor cells, compared to MOs. This increased complexity allows for the identification of cell-type-specific phenotypes, particularly those involved in cross-lineage interactions that are not apparent in neuron-centric models.

Our transcriptomic data revealed a transcriptionally distinct dopaminergic neuron cell cluster in PD MGL-MOVOs, the DNs2 cluster, which was absent in the HC group. Neurons in the DNs2 cluster were characterized by a downregulation of genes related to synaptic transmission, synaptic adhesion and scaffolding, and axonal guidance, pointing towards reduced functional integration. Alterations in synaptic transmission and activity have been reported in genetic PD models ^63,67^. However, despite the identification of vulnerable dopaminergic neuron subpopulations in human post-mortem PD samples ^68^, no dopaminergic sub-cluster had yet been linked to a lower synaptic functionality. Importantly, DNs2 also showed increased expression of genes associated with cell death pathways, including *CASP7* ^69^ and *GPX4* ^70,71^, suggesting that these neurons may be in a vulnerable or pre-degenerative state ^65^. The identification of DNs2 neurons opens the door to future studies of dopaminergic neuron vulnerability in midbrain assembloid models of PD. In addition, the MGL-MOVO’s multicellular architecture enables the discovery of complex, also non-neuronal phenotypes, providing insights that may be missed in neuron-centric models.

MEA results further indicate an impaired neuronal activity in PD MGL-MOVOs, consistent with a prior study on a LRRK2-G2019S cerebral organoid model ^72^. In parallel, microglia displayed reduced ramification, which correlates with immunostainings of postmortem human samples of idiopathic PD patients revealing a more amoeboid shape of microglia cells ^73^. MGL-MOVOs from the PD group showed a higher proportion of cells expressing the pro-inflammatory cytokines TNFα, IL1β, IFNγ, and IL6 compared to HC. In line with these results, transcriptomic human post-mortem samples from idiopathic PD revealed an overexpression of *IL1B* by microglia cells, indicating a pro-inflammatory activation ^73^. Importantly, *TREM2*, a key regulator of microglial homeostasis, phagocytosis, and immune modulation, was expressed at higher levels in HC MGL-MOVOs compared to PD MGL-MOVOs. TREM2’s role in regulating phagocytosis, supressing excessive inflammatory responses, and maintaining microglial fitness is well established ^24–26^. Its reduced expression in PD MGL-MOVOs suggests impaired microglial regulatory capacity, consistent with the observed shift toward a pro-inflammatory phenotype. Given TREM2’s emerging role as a therapeutic target in neurogenerative diseases ^74^, MGL-MOVOs could serve as a valuable platform for studying TREM2-associated signaling and its modulation in disease context. Overall, these results demonstrate that MGL-MOVOs are a suitable model for PD, and they recapitulate phenotypes observed in other cell culture systems, as well as in human post-mortem samples.

These results show that MGL-MOVOs provide a powerful and physiological relevant *in vitro* model for studying midbrain-specific neurodegeneration, The ability to recapitulate neurovascular interactions, identify disease-associated dopaminergic subtypes, and capture inflammatory dysregulation offers unique advantages for mechanistic studies of Parkinson’s disease, particularly those involving glial-neuronal-vascular crosstalk.

Beyond disease modeling, the scalable and reproducible nature of MGL-MOVOs makes them well-suited for drug screening, target validation, and personalized medicine applications. Their multi-lineage complexity allows assessment of compound effects not only on neurons but also on vascular and immune compartments, better mimicking *in vivo* drug responses and toxicity profiles.

## Methods

### 2D cell culture

Generation of iPSCs (Table 1) was done as described in ^75^. In short, 6-well plates (Thermo Scientific, 140675) were coated with Matrigel^®^ (Corning, 354277), and 300K cells were plated per well. We used Essential 8 Basal medium (Thermo Scientific, A1517001) for cell maintenance, with a daily medium change. The first 24 h after cell splitting or thawing, the medium was supplemented with 10 µM ROCK Inhibitor (Y-27632, Millipore, SCM075). When confluence reached 70-90%, cells were split by incubating for 5 minutes with Accutase^®^ (Sigma-Aldrich, A6964). The generation of neural progenitor cells from iPSCs and their maintenance were performed as described in ^8,19^. The derivation of macrophage precursors from iPSCs and their maintenance were performed as described in ^76^.

### 3D cell culture

#### Midbrain organoid generation and culture

We generated and cultured midbrain organoids as described in ^8,19^. In short, 6,000 cells were seeded per well in an Ultra-Low Attachment 96-well plate (Merck, CLS3474) and cultured in maintenance medium (N2B27 medium supplemented with 0.2 mM Ascorbic acid (Sigma-Aldrich, A4544), 3 μM CHIR 99021 (Axon Medchem, CT 99021), 0.5 μM Smoothened Agonist, SAG (Merk Millipore, 566660), 2.5 μM SB-431542 (Abcam, ab120163), 0.1 μM LDN-193189 (Sigma-Aldrich, SML0559)) for 2 days. Then, the medium was replaced to induce ventral midbrain patterning (day 0 of dopaminergic differentiation) by removing SB and LDN. Two days after, CHIR concentration was reduced to 0.7 μM. On day 6 of dopaminergic differentiation, the medium was replaced by maturation medium (N2B27 plus 0.2 mM Ascorbic acid, 10ng/mL Brain Derived Neurotrophic Factor, BDNF (Peprotech, 450-02), 10 ng/mL Glial-Derived Neurotrophic Factor, GDNF (Peprotech, 450-10), 1 pg/mL TGF-β3 (Peprotech, 100-36E), 10 μM DAPT (R&D Systems, 2634/10) and 2.5 ng/mL Activin A (Thermo Scientific, PHC9564)). We kept the spheroids in static conditions (without shaking) until day 6 (day 4 of dopaminergic differentiation - when they were cocultured with macrophage precursors - or day 8 (day 6 of dopaminergic differentiation), when they were embedded in Matrigel^®^ (Corning, 354277) as described in ^7^ (Monzel et al., 2017c) or co-cultured with vascular organoids.

#### Vascular organoid generation and culture

We generated and cultured vascular organoids as described in ^18^ (Wimmer et al., 2019b) with slight modifications. In short, 6000 iPSCs were seeded per well in a 96-well plate resuspended in vascular differentiation medium (DMEM/F12 medium (Thermo Fisher, 21331020), 20% KOSR (Thermo Fisher, 10828028), 1x Glutamax (Thermo Fisher, 35050061), 1x NEAA (Thermo Fisher, 11140050)), supplemented with 50 μM ROCK Inhibitor (Y-27632, Millipore, SCM075) and placed in an incubator with hypoxic conditions (5% O_2_). On day 3 after seeding, the medium was exchanged and supplemented with 12 μM CHIR99021. On days 5, 7 and 9, the medium was exchanged again, and supplemented with 30 ng/mL BMP4 (Peprotech, 120-05), 30 ng/mL VEGF-A (Peprotech, 100-20) and 30 ng/mL FGF-2 (Peprotech, 100-18B). On day 11, the exchanged medium contained 30 ng/mL VEGF-A, 30 ng/mL FGF-2 and 10 μM SB-43152. On day 13, vascular organoids were used for co-culture with midbrain organoids and embedded in Matrigel^®^ as described in ^7^ (Monzel et al., 2017c) (Figure 1A).

#### Co-culture of midbrain organoids with vascular organoids

Co-culture of midbrain and vascular organoids was done by co-embedding at day 8 of midbrain differentiation and day 13 of vascular organoid culture (Figure 1A). Midbrain and vascular organoids were placed together in a Matrigel^®^ droplet of around 30 µl and let polymerize for 25 minutes at 37°C. After that, they were transferred into 24-well plates and cultured with midbrain maturation medium on an orbital shaker rotating at 80 rpm.

#### Co-culture of midbrain organoids and midbrain-vascular assembloids with macrophage precursors

Two days before the embedding, at day 6 (day 4 of dopaminergic differentiation), midbrain spheroids were organized into four sub-groups: MOs, MGL-MOs, MOVOs, and MGL-MOVOs (Figure 2A). The culture medium of the four groups was exchanged into co-culture medium ^11^ (Advanced DMEM/F12 (Thermo Fisher, 12634010), 1x N2 (Thermo Fisher, 17502001), 1x Pen/Strep (Invitrogen, 15140122), 1x Glutamax (Thermo Fisher, 35050061), 50 µM 2-mercaptoethanol (Thermo Fisher, 31350-010), 100 ng/mL IL-34 (Peprotech, 200-34), 10 ng/mL GM-CSF (Peprotech, 300-03), 10 ng/mL BDNF, 10 ng/mL GDNF, 10 μM DAPT and 2.5 ng/mL Activin A. For MGL-MOs and MGL-MOVOs, macrophage precursors were resuspended in culture medium and added on the well (186K macrophage precursors per MO and 450K macrophage precursors per MOVO). Two days later, we proceeded with the embedding as described previously.

#### Immunofluorescence staining in 3D

At day 30 of differentiation, organoids and assembloids were fixed with 4 % Formaldehyde overnight (16h) at 4°C and washed 3 times with PBS for 15 min. For immunostainings of sections, they were embedded in 4 % low-melting point agarose (Biozym, Cat. No. 840100) in PBS. A vibratome (Leica VT1000 S) was used to slice the organoids and assembloids into 70 or 120 μm sections. Blocking and permeabilization of sections was done with 0.5% Triton X-100, 0.1% sodium azide, 0.1% sodium citrate, 2% BSA, and 5% donkey serum in PBS for 90 min at room temperature. Primary antibodies (Table 2) were diluted in 0.1% Triton X-100, 0.1% sodium azide, 0.1% sodium citrate, 2% BSA, and 5% donkey serum, and sections were incubated with the primary antibody solution for 48 h at 4°C in a shaker. Then, sections were washed 3 times with PBS and incubated with secondary antibodies in 0.05% Tween-20 in PBS for 2 h at RT. After that, they were washed 3 times with 0.05% Tween-20 in PBS and one time with water. Sections were mounted in Fluoromount-G mounting medium on a glass slide.

#### Imaging

We used a Zeiss LSM 710 confocal laser-scanning microscope for representative images and the ZEN blue Software for adjustments and modifications.

#### Image analysis

For image analysis of IBA1 and CD31 morphology, we used the Imaris software (Oxford Instruments). We used the ‘Filaments’ tracer to segment positive cells, detect and modify seeding points, and trace filaments and processes. Statistics were run with data per cell in GraphPad Prism.

#### Hypoxyprobe assay

The Hypoxyprobe™ Omni Kit (Hypoxyprobe, HP3-200Kit) was used to measure hypoxia levels in MOs and MOVOs. Pimonidazole was diluted in media at a concentration of 250 µM, and MOs and MOVOs were incubated in pimonidazole solution for 2h at 37°C. Then, fixation was performed with 4% Formaldehyde overnight at room temperature, followed by agarose embedding and sectioning into 70 µm sections. The staining was done as described in “Immunofluorescence staining in 3D”, using the PAb2627AP antibody against pimonidazole and a secondary HRP-conjugated antibody. Diaminobenzidine (DAB) tablets were used to proceed with the staining (SIGMAFAST™ 3,3′-Diaminobenzidine tablets, Sigma-Aldrich, D4293). A DAB tablet and a Urea hydrogen peroxide tablet were mixed in 5 ml water. The solution was applied to MO and MOVO sections for < 1 min until the colour developed visibly. Then, sections were washed with water and mounted with Fluoromount-G mounting medium on a glass slide. Images were acquired in a stereomicroscope (Nikon, SMZ25-SMZ18) and positive areas were quantified with ImageJ.

#### LDH assay

For the LDH viability assay, we used supernatant pooled from 3 MOs and MOVOs per batch, and 3 batches were considered. We used the line HC1 for this assay. We snap-froze supernatants the day of collection, and kept them at -80°C until the day the assay was performed. The LDH-Glo™ Cytotoxicity Assay (Promega, J2380 and J2381) kit was used, and samples were loaded into 96-well plates with opaque walls. Reagents were thawed at RT, and samples were thawed on ice. We used 25 µl of supernatant and 25 µl of LDH detection reagent per well and incubated for 60 min at RT. Luminescence was recorded in a TECAN reader. We normalized the OD values to organoid and assembloid size (area measured in ImageJ). After assessing the normality of the data, statistics were run using a t-test in GraphPad Prism.

### Single-nuclei RNA sequencing (snRNA-Seq)

#### Sample pre-processing

MOs and MGL-MOVOs from the line HC1 were used for this protocol. We collected 3 MOs or MGL-MOVOs per batch and 3 batches. At day 30 of differentiation, samples were snap-frozen and stored at -80°C. For the nuclei extraction, MOs or MGL-MOVOs from the 3 batches were pooled, resulting in two sample groups.

#### Nuclei extraction and library preparation

To extract nuclei, frozen MOs and MGL-MOVOs were transferred into a 1.5 ml vial with 0.5ml ice cold cell lysis buffer containing 10 mM Tris-HCl, pH 7.4, 10 mM NaCl, 3 mM MgCl2, 0.1% IGEPAL CA-630, 1% SUPERase In RNase Inhibitor (20 U/µL, Ambion) and 2% BSA and physically disrupted using a disposable pistille. The mixture was filtered through a 10um filter and the nuclei were centrifuged down at 500g for 5min at 4°C. The supernatant was discarded, and the nuclei were resuspended in 0.5 ml nuclei suspension buffer containing 10 mM Tris-HCl, pH 7.4, 10 mM NaCl, 3 mM MgCl2, 1% SUPERase In RNase Inhibitor (20 U/µL, Ambion) and 2% BSA. Nuclei were counted via Neubauer chamber and Trypan Blue staining, centrifuged down at 500g for 5min at 4°C, the supernatant was discarded, and the nuclei were resuspended in the appropriate amount of nuclei suspension buffer to yield a single nuclei solution of ∼1000 nuclei/ul. The single nuclei suspensions were processed into labelled cDNA and subsequently NGS libraries following manufacturer’s instructions using Next GEM Single Cell 3’ (version 3.1) chemistry for dual indexing on a Chromium device (10X Genomics). cDNA and libraries were quantified using a HS dsDNA assays on a Qubit 2 instrument (Thermo fisher) and fragment size distribution was determined via High Sensitivity DNA assay on a Bioanalyzer2100 instrument (Agilent). Final library molarity was determined via NEBNext Library Quant Kit for Illumina (NEB) on a QuantStudio 5 Real-Time PCR System (Applied Biosystems). Libraries were equimolarly pooled, clustered at 1000pM on a P3 flowcell and sequenced PE R1 28 cycles 2*10 index cycles and R2 90 cycles using 100 cycle SBS chemistry on a NextSeq2000 instrument (Illumina). Data was demultiplexed, converted to fastq format and aligned to reference genome hg19 using Cellranger (version 5.0.1) (10X Genomics).

#### Data analysis

Nuclei expressing less than 500 genes were filtered out. From those, only nuclei with at least 750 total transcript counts but not more than 25,000 total transcript counts were retained for the analysis. In addition, nuclei with mitochondrial or ribosomal gene expression greater than 5% were removed. Downstream data analysis was done using Seurat package ^77^ (version 5.2.1) on R (version 4.0.2). All samples were integrated into a single Seurat object following Seurat integration workflow ^77^ based on the 2500 most variable genes and 50 principal components. Cellular clusters were identified, applying the Louvain algorithm modularity optimization with a resolution of 0.1. Visualization via Uniform Manifold Approximation and Projection (UMAP) ^78^ revealed in total of 13 distinct cell types. Cell types were identified using a binarized marker gene list of embryonic midbrain cell types given by La Manno et al. ^79^ . Cell identity detection was based on the amount of marker genes from embryonic midbrain cell populations, expressed in every cell cluster of our dataset. Cell cluster identity was further refined using PangloaDB-human single-cell RNA sequencing experiment database ^80^ and GeneAnalystics online tool ^81^. In the latter, marker genes of each cell cluster found using FindAllMarkers function of Seurat, were given in the tool for cell type identification choosing in vitro parameter for brain cells. Differentially expressed genes (DEGs) were detected using the FindMarkers function of the Seurat using the default Wilcoxon Rank Sum test, considering only genes expressed in at least 10% of cells in either of the two conditions compared. Significantly differentially expressed genes were considered with p.adjust<0.05. Pathway enrichment analysis was done using MetaCore (version 21.4, Clarivate).

### Multi-electrode array

To prepare 48-well MEA plates (Axion, M768-tMEA-48B-5), wells were coated with 0.1 mg/ml poly-D-lysine (Sigma-Aldrich, P7886) and incubated at 37 °C, 5 % CO_2_ for 1 h, followed by an overnight coating with 10 μg/ml laminin (BioLamina, LN111). On day 7 of co-culture, MGL-MOVOs were placed in the centre of the well, directly over the electrodes, and incubated for 5 min to promote media evaporation and adherence to the well surface. Then, 25 µl Geltrex was added on top of each MGL-MOVO to prevent detachment, followed by a 15-minute incubation to allow polymerization. Afterward, 300 μl fresh media was added to each well, and plates were maintained under static conditions at 37 °C, 5 % CO_2_. Spontaneous extracellular field potentials were recorded using the Axion Maestro 768-channel MEA system and the AxIS software (Axion BioSystems, v.2.1.) and analysed with R (version 4.3.0). Assembloid activity was analysed using data from active electrodes. Electrodes that recorded no activity or fewer than three spikes within five minutes were excluded from the analysis. Statistical significance was assessed using Wilcoxon signed-rank test. Outliers were identified and excluded based on the 1.5 interquartile range (IQR) method, calculated from the 25^th^ and 75^th^ percentiles.

### Flow cytometry

For flow cytometry analysis, four embedded MGL-MOVOs from each of the HC1, HC4, HC5, and PD cell lines were pooled and digested in papain solution (0.18 % Papain, Sigma-Aldrich; 0.04 % EDTA, Sigma-Aldrich; 0.04% L-Cysteine, Sigma-Aldrich; in DMEM/F12) for 50 min at 37 °C. After digestion, papain was gently aspirated, and MGL-MOVOs were dissociated at 37 °C using Accutase. Gentle pipetting was used to further aid dissociation and ensure the formation of a single-cell suspension. To halt the digestion, PBS containing 0.5 % BSA and 0.5 % Trypsin inhibitor (Roche, 10109878001) was added, and the cells were transferred into Eppendorf tubes for centrifugation at 500 x g for 2 min. Following centrifugation, cell pellets were washed once with PBS and resuspended in DMEM/F12.

Microglia were identified as CD11b^+^. For intracellular staining of cytokines, cells were washed once with FACS buffer (1 % FBS and 5 mM EDTA, pH 8.0, in 1 x PBS), then fixed and permeabilised using the BD Pharmingen^TM^ Transcription Factor Buffer Set (BD Biosciences, 562574). Permeabilised cells were stained for 2 hours at RT in the dark with antibodies (Table 2) and Zombie Green^TM^ Fixable Viability Kit (Biolegend, 423112) in 1 x Perm/Was Buffer. After incubation, cells were washed twice with 1 x Perm/Wash Buffer before flow cytometry analysis. Data were acquired on a BD LSRFortessa Cell analyzer (BD Biosciences) and analysed using FlowJo v10 software (Tree Star).

### Statistical analysis

Unless specified otherwise, statistical analyses and data visualization were performed using GraphPad Prism (version 10.2.3) or R software (version 4.4.3). The Shapiro-Wilk test was applied to assess the normality of the data distribution, and potential outliers were identified and excluded using the ROUT method with a false discovery rate (Q) of 1%. For comparison between two groups, the Mann-Whitney U test was used when data did not meet normality assumption, while an unpaired, two-tailed t-test was employed for normally distributed data. Significance asterisk represent *p < 0.05, **p<0.01, ***p<0.001, ****p<0.0001. Error bars represent mean + standard derivation (SD).

## Data availability

All original and processed data, along with the metadata supporting the findings of this study, are publicly available at: DOI: 10.17881/qaxz-gp42. Gene expression datasets have been deposited in the Gene Expression Omnibus (GEO) under the accession number GSE280501.

All scripts used for preprocessing, analysis, and visualization are available at: https://gitlab.com/uniluxembourg/lcsb/developmental-and-cellular-biology/zimmermann_2025.

## Acknowledgments

The authors thank Dr. Jared Sterneckert, as well as Dr. Nico J. Diederich and Laura Longhino from Centre Hospitalier de Luxembourg for providing iPSC lines. CS was supported by Fonds National de la Recherche Luxembourg (C19/BM/13535609). The JCS lab is supported by the Fonds National de la Recherche (FNR) Luxembourg (PRIDE17/12244779/PARK-QC; PRIDE21/16749720/NextImmune2 ; FNR/PoC16/11559169, FNR/ NCER13/BM/11264123). We also thank the private donors who support our work at the Luxembourg Centre for Systems Biomedicine.

## Rights Retention Statement

This research was funded in whole by FNR Luxembourg. For the purpose of Open Access, the author has applied a CC BY public copyright license to any Author Accepted Manuscript (AAM) version arising from this submission.

## Author contributions

SSS, ASZ, and CS designed and conducted the experiments, interpreted the data, and drafted the manuscript. AZ and KB analysed the data. KH conducted specific experiments. SR and LF contributed with knowledge and practical information. CS, MS, and JCS coordinated and conceptualized the study. All authors reviewed and approved the final manuscript and agreed to be accountable for their contributions.

## Competing interests

JCS is co-founder and shareholder of the biotech company Organo Therapeutics SARL. This company used midbrain organoids and assembloids for in vitro disease modeling and drug discovery.

## Materials and Correspondence

For correspondence and material requests, please contact Jens Schwamborn at.

## Ethics Statement

Written informed consent was obtained from all individuals who donated samples to this study, and all work with human stem cells was done after approval of the national ethics board, Comité National d’Ethique de Recherche (CNER), under the approval numbers 201305/04 and 201901/01.

## Notes

https://doi.org/10.17881/qaxz-gp42

