## Supplementary figures and images for "Vascularized midbrain assembloids show neuroinflammation and dopaminergic neuron vulnerability in Parkinson’s Disease"

### Supplementary Figures

##### Supplementary Figure 1

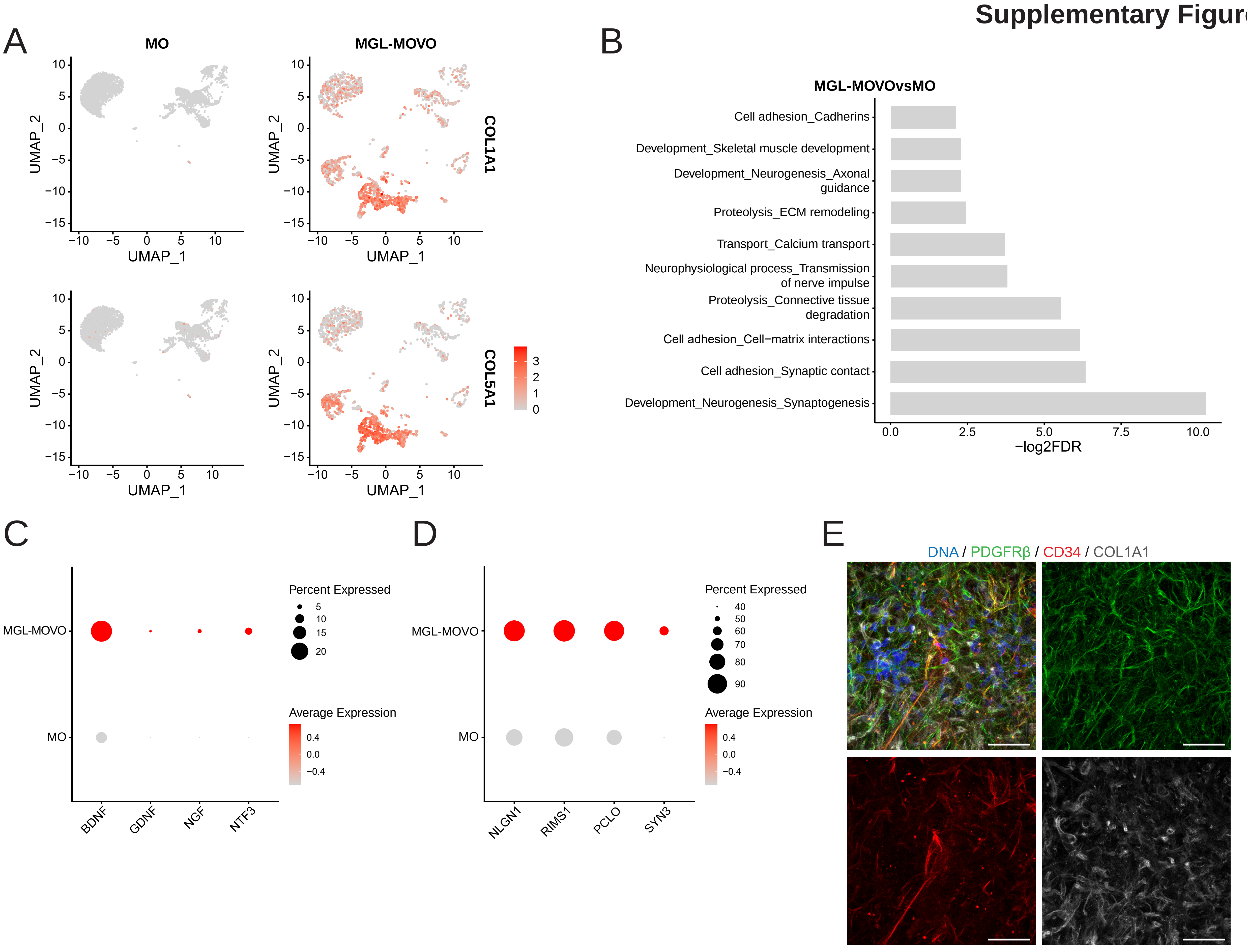

#### Supplementary Figure 2

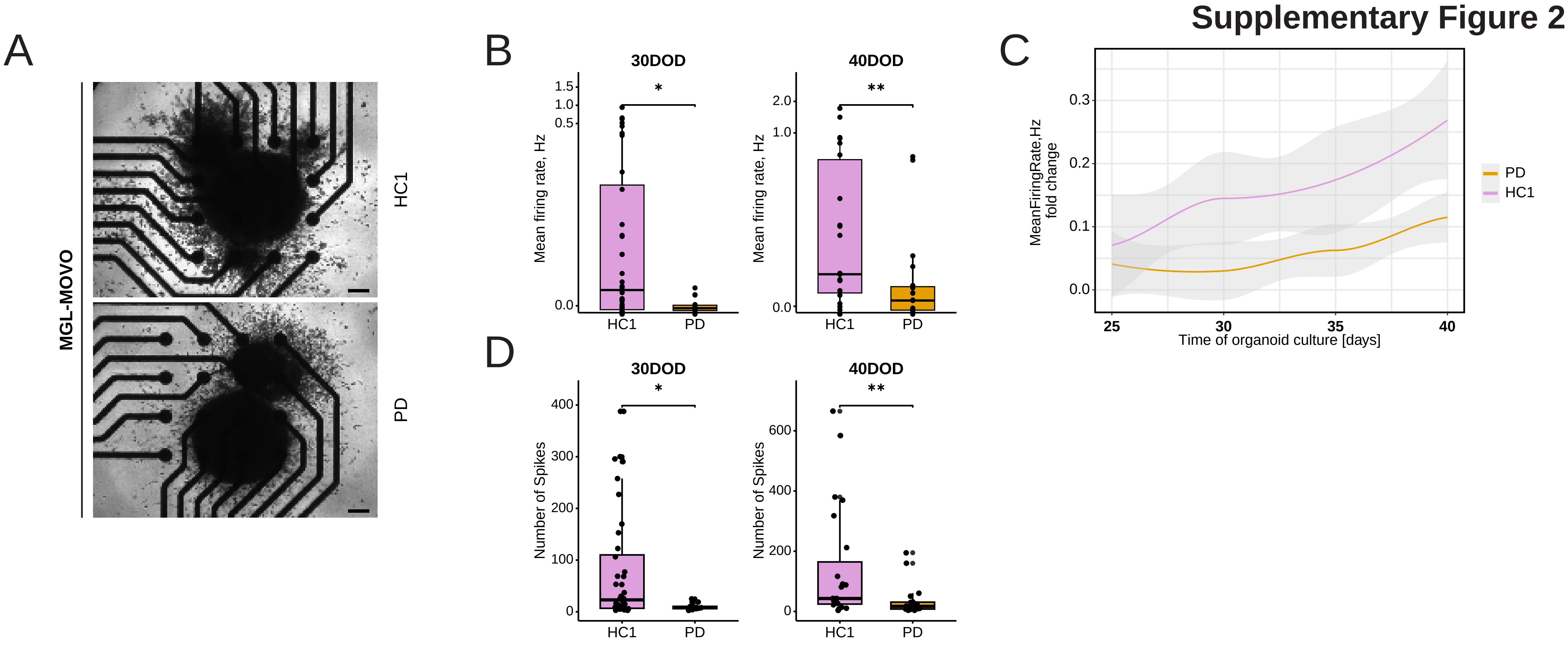
