## Supplementary Table 1 for "Vascularized midbrain assembloids show neuroinflammation and dopaminergic neuron vulnerability in Parkinson’s Disease"

**Supplementary Table 1 related to experimental procedures**. iPS cell lines used in this study.

| Sample ID | Diagnosis | Genotype | Sex | Age of sampling | Source |
| --- | --- | --- | --- | --- | --- |
| HC1 | HC | WT | F | Cord Blood | GIBCO/A13777 |
| HC2 | HC | WT | F | 81 | Reinhardt et al., 2013 |
| HC3 | HC | WT | M | 55-59 | Qing et al., 2017 |
| HC4 | HC | WT | F | 65 | Dr. Nico J. Diederich (Centre Hospitalier de Luxembourg) |
| HC5 | HC | WT | F | 63 | Dr. Nico J. Diederich (Centre Hospitalier de Luxembourg) |
| PD | PD | LRRK2 G2019S | F | 81 | Reinhardt et al., 2013 |
