## Supplementary Table 2 for "Vascularized midbrain assembloids show neuroinflammation and dopaminergic neuron vulnerability in Parkinson’s Disease"

**Supplementary Table 2 related to experimental procedures**. Antibodies used in this study. IF = Immunofluorescence.

| Antibody | RRID | Host species | Source | Ref.-No. | Assay | Dilution |
| --- | --- | --- | --- | --- | --- | --- |
| TH | *AB_297840* | Rabbit | Abcam | ab112 | 3D IF | 1:1000 |
| TUJ1 | *AB_570918* | Chicken | Millipore | AB9354 | 3D IF | 1:1000 |
| CD31 | *AB_2114471* | Mouse | DAKO | M082301 | 3D IF | 1:200 |
| CD31 | *AB_726362* | Rabbit | Abcam | ab28364 | 3D IF | 1:100 |
| CD34 | *AB_2063006* | Mouse | DAKO | M7165 | 3D IF | 1:100 |
| MAP2 | *AB_2138153* | Chicken | Abcam | ab5392 | 3D IF | 1:1000 |
| Collagen IV | *AB_305584* | Rabbit | Abcam | ab6586 | 3D IF | 1:200 |
| Collagen I | *AB_731684* | Rabbit | Abcam | ab34710 | 3D IF | 1:500 |
| α-SMA | *AB_2223021* | Rabbit | Abcam | ab5694 | 3D IF | 1:100 |
| VE-Cadherin | *AB_870662* | Rabbit | Abcam | ab33168 | 3D IF | 1:300 |
| Aquaporin 4 | *AB_626695* | Mouse | Santa Cruz | sc-32739 | 3D IF | 1:100 |
| Occludin | *AB_881773* | Rabbit | Abcam | ab31721 | 3D IF | 1:100 |
| PDGFRβ | *AB_355339* | Goat | R&D systems | AF385-SP | 3D IF | 1:100 |
| GFAP | *AB_177521* | Chicken | Millipore | AB5541 | 3D IF | 1:1000 |
| PAb2627AP | *-* | Rabbit | Hypoxyprobe | PAb2627AP | Hypoxyprobe | 1:100 |
| Anti-Rabbit HRP | *AB_772206* | Donkey | GE Healthcare | NA934 | Hypoxyprobe | 1:1000 |
| IBA1 | *AB_2224402* | Goat | Abcam | ab5076 | 2D/3D IF | 1:250 |
| Alexa Fluor® 647 Anti-chicken | *AB_2340379* | Donkey | Jackson Immuno | 703-605-155 | 3D IF | 1:1000 |
| Alexa Fluor® 488 anti-goat | *AB_2534102* | Donkey | Invitrogen | A-11055 | 3D IF | 1:1000 |
| Alexa Fluor® 568 anti-goat | *AB_2534104* | Donkey | Thermo Fisher | a11057 | 3D IF | 1:1000 |
| Alexa Fluor® 647 anti-goat | *AB_2535864* | Donkey | Invitrogen | a21447 | 3D IF | 1:1000 |
| Alexa Fluor® 488 anti-rabbit | *AB_2535792* | Donkey | Thermo Fisher | a21206 | 2D/3D IF | 1:1000 |
| Alexa Fluor® 568 anti-rabbit | *AB_2534017* | Donkey | Invitrogen | a10042 | 2D/3D IF | 1:1000 |
| Alexa Fluor® 647 anti-rabbit | *AB_2536183* | Donkey | Invitrogen | a31573 | 2D/3D IF | 1:1000 |
| Alexa Fluor® 488 anti-mouse | *AB_141607* | Donkey | Invitrogen | a21202 | 2D/3D IF | 1:1000 |
| Alexa Fluor® 568 anti-mouse | *AB_2534013* | Donkey | Invitrogen | a10037 | 2D/3D IF | 1:1000 |
| Hoechst 33342 Solution (20 mM) | *-* | - | Invitrogen | 62249 | 2D/3D IF | 1:10000 |
| Alexa Fluor® 647 anti-human IL-1β | *AB_2124352* | Mouse | Biolegend | 511707 | Flow cytometry | 1:500 |
| APC/Cyanine7 anti-human TNF-α | *AB_2562870* | Mouse | Biolegend | 502944 | Flow cytometry | 1:500 |
| Brilliant Violet 605^TM^ anti-human CD11b | *AB_2562021* | Mouse | Biolegend | 301332 | Flow cytometry | 1:500 |
| PE/Cyanine7 anti-human IL-6 | *AB_2572041* | Rat | Biolegend | 501119 | Flow cytometry | 1:500 |
| PerCP/Cyanine5.5 anti-human IFN-γ | *AB_2566187* | Mouse | Biolegend | 506528 | Flow cytometry | 1:500 |
| Zombie Green^TM^ Fixable Viability Kit | *-* | - | Biolegend | 423112 | Flow cytometry | 1:1000 |
