## Supplementary Table 3 for "Vascularized midbrain assembloids show neuroinflammation and dopaminergic neuron vulnerability in Parkinson’s Disease"

**Supplementary Table 3 related to snRNA-seq**. Percentage of cell identities in HC1 MOs and HC1 MGL-MOVOs.

| Cell cluster | MO [%] | MGL-MOVO [%] |
| --- | --- | --- |
| Radial glia | 47.56 | 18.74 |
| Neurons | 33.64 | 6.55 |
| Endothelial cells 1 | 0 | 9.04 |
| Mesoderm-like cells, progenitors | 0 | 31.76 |
| Cholinergic neurons | 7.52 | 3.18 |
| Megakaryocyte precursor cells | 0 | 19.58 |
| Pericytes | 0.35 | 0.19 |
| Astrocytes | 0.12 | 0.08 |
| Endothelial cells 2 | 0 | 3.18 |
| Dopaminergic neurons 1 | 10.80 | 2.41 |
| Microglia | 0 | 3.72 |
| Endothelial cells 3 | 0 | 1.57 |
