## Supplementary Table 4 for "Vascularized midbrain assembloids show neuroinflammation and dopaminergic neuron vulnerability in Parkinson’s Disease"

**Supplementary Table 4 related to snRNA-seq**. Percentage of cell identities in HC1 and PD MGL-MOVOs.

| Cell cluster | HC1 [%] | PD [%] |
| --- | --- | --- |
| Radial glia | 18.74 | 29.31 |
| Neurons | 6.55 | 8.18 |
| Endothelial cells 1 | 9.04 | 10.26 |
| Mesoderm-like cells, progenitors | 31.76 | 15.64 |
| Cholinergic neurons | 3.18 | 3.64 |
| Megakaryocyte precursor cells | 19.58 | 14.31 |
| Pericytes | 0.19 | 5.76 |
| Astrocytes | 0.08 | 4.68 |
| Dopaminergic neurons 2 | 0.00 | 1.48 |
| Endothelial cells 2 | 3.18 | 3.76 |
| Dopaminergic neurons 1 | 2.41 | 0.81 |
| Microglia | 3.72 | 0.88 |
| Endothelial cells 3 | 1.57 | 1.27 |
